# An actin-like filament from *Clostridium botulinum* exhibits a novel mechanism of filament dynamics

**DOI:** 10.1101/2022.03.07.483215

**Authors:** Adrian Koh, Samson Ali, David Popp, Kotaro Tanaka, Yoshihito Kitaoku, Naoyuki Miyazaki, Kenji Iwasaki, Kaoru Mitsuoka, Robert C. Robinson, Akihiro Narita

## Abstract

Here, we report the discovery of a ParM protein from *Clostridium botulinum* (CBgs-ParM), which forms a double-stranded polar filament. CBgs-ParM shares many similarities in its basic filament architecture with actin, however, Pi release after nucleotide hydrolysis induces a large lateral strand shift of ~2.5 nm. We identified the ParR (CBgs-ParR) that acts as a nucleation factor in the initial stage of polymerization, similar to ParR from *Escherichia coli* plasmid R1. CBgs-ParR also functions as a depolymerization factor, probably by recognizing the structural change in the CBgs-ParM filament after Pi release. Comparison with CBH-ParM, another ParM from *Clostridum botulinum*, showed that subunit-subunit interacting regions largely differ, preventing co-polymerization, implying a selection pressure in evolution to prevent interference between different ParMRC systems.

## Introduction

While high-copy number plasmids can be propagated in bacteria by passive diffusion for successful inheritance from mother to daughter cells, low-copy number plasmids require an active transport system to ensure their proper inheritance. ParM, an actin homolog in prokaryotes, is a member of the ParMRC system (Bharat *et al*, 2015; Salje *et al*, 2010), which contributes to plasmid segregation in combination with ParR, another protein, and *parC*, a DNA sequence on the target plasmid. Each ParM unit binds one nucleotide, ATP or GTP, and forms a filament via polymerization, which pushes the plasmids apart. The bound ATP or GTP is hydrolyzed in the filament after polymerization. The hydrolysis of ATP or GTP and subsequent release of inorganic phosphate (Pi) largely destabilizes the filament and induces depolymerization (Garner *et al*, 2004; Jiang *et al*, 2016; Koh *et al*, 2019), a critical step for filament turnover. ParR is considered to have several properties: as an adapter that connects ParM to *parC;* as a nucleator that initiates ParM polymerization; and as a stabilizer of the filament (Garner *et al*, 2007; Gayathri *et al*, 2012).

However, ParMRC systems are highly divergent (Bharat *et al.*, 2015; Jiang *et al.*, 2016; Koh *et al.*, 2019; Popp *et al*, 2012) and a wide range of biochemical and structural features of the constituent molecules, in different bacteria, remain to be elucidated. Here, we report a novel ParM, CBgs-ParM, from *Clostridium botulinum*, which shows several unique features.

## Results

### CBgs-ParMRC

CBgs-ParM (ED787363.1) and CBgs-ParR (EDT87283.1) sequences were identified through homology searches using BLAST on a whole genome shotgun sequence of *Clostridium botulinum* Bf (Accession ID: ABDP01000001). “gs” represents “genome shotgun”. A putative *parC* sequence containing palindromic sequence repeats was also found immediately upstream of CBgs-ParMR (Fig. 1A). The putative *parC* sequence bound to ParR in an electrophoretic mobility shift assay (Fig. 1B), indicating that the putative sequence acts as the CBgs-*parC*.

**Figure 1:**
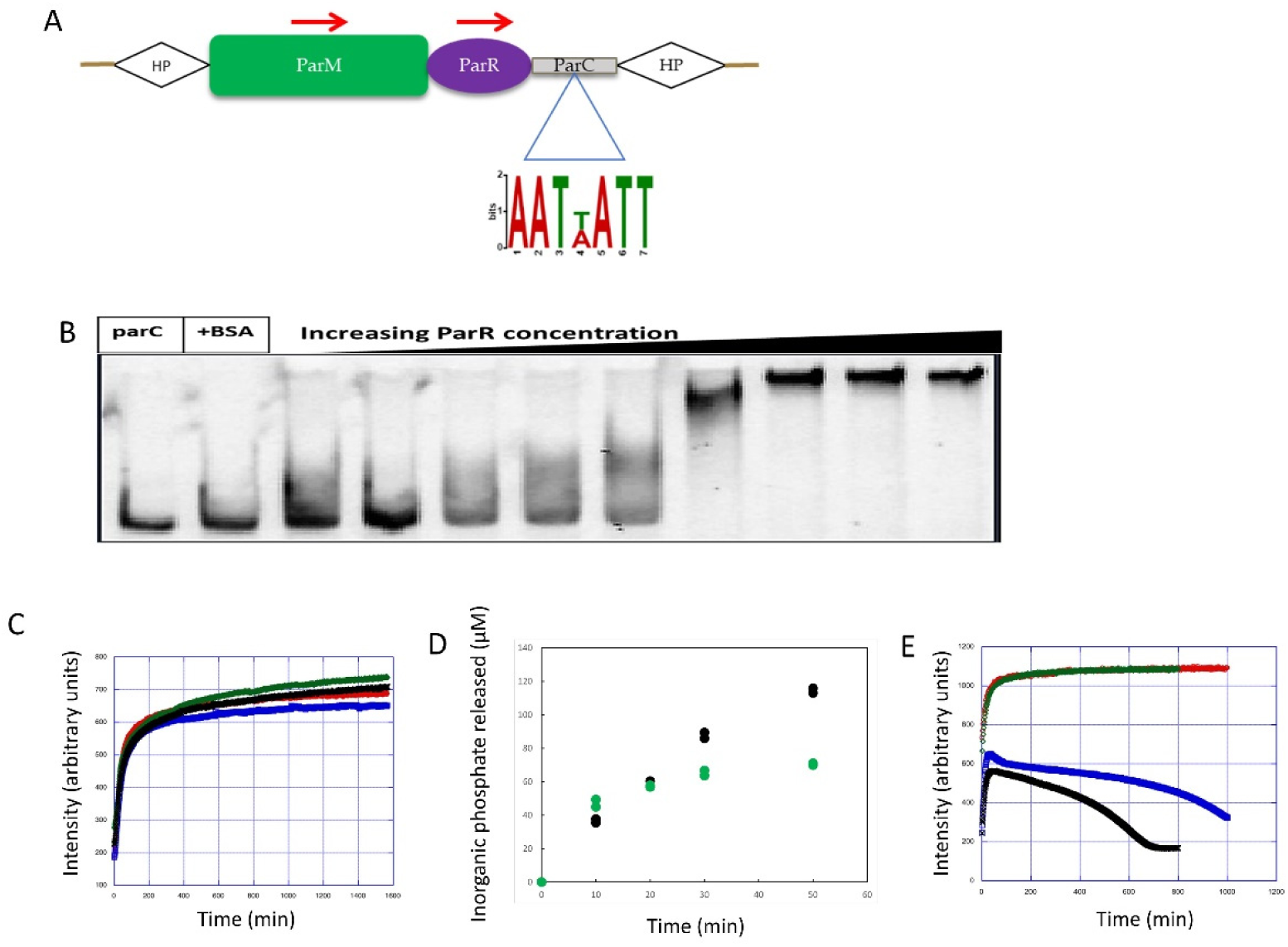
Characterization of the CBgs-ParMRC system. A ParMRC system present on the genome of *C. botulinum* strain Bf. CBgs-*parC* includes three palindromic repeats. HP represents hypothetical proteins on both sides of the *parMRC* operon. B Electrophoretic mobility shift assay of CBgs-*parC* with 10× to 1000× molar excess of ParR. C Light scattering time courses of CBgs-ParM polymerization. 10 μM CBgs-ParM green, ATP; black, GTP; red, ADP; blue, GDP. D Pi release from 21 μM CBgs-ParM, black with GTP and green with ATP. Two measurements for each nucleotide are superposed. E Light scattering time courses of CBgs-ParM polymerization in the presence of CBgs-ParR, blue, with ATP; black, with GTP; Corresponding time courses without CBgs-ParR, red, with ATP; green, with GTP.

### Polymerization assay and ParR-ParM interactions

CBgs-ParM polymerized with GTP, ATP, GDP and ADP at similar rates as judged by an increase in light scattering over time (Fig. 1C). After polymerization, CBgs-ParM remained as a filament without undergoing bulk depolymerization judged by constant light scattering intensity (Fig. 1C), unlike previously studied ParMs(Jiang *et al.*, 2016; Koh *et al.*, 2019). Supernatant CBgs-ParM concentrations in a sedimentation assay indicated that the critical concentration for polymerization were similar for the different nucleotides, determined to be 2.3 ± 0.1 μM, 2.8 ± 0.1 μM, 1.2 ± 0.1 μM, and 3.6 ± 0.4 μM with ATP, ADP, GTP, and GDP, respectively (Fig. S1). The critical concentration dependence of CBgs-ParM on the nucleotide state was considerably smaller than that of actin (Fujiwara *et al*, 2007) or *E. coli* ParM-R1 (Garner *et al.*, 2004). Continuous phosphate release beyond the concentration of the protein (21 μM CBgs-ParM) was observed (Fig. 1D), suggesting subunit exchange from the ends of the filament after the initial polymerization, consistent with a process such as treadmilling (Narita, 2011; Pollard & Borisy, 2003; Wegner, 1976).

In the presence of CBgs-ParR, the CBgs-ParM initial polymerization rate was increased, indicating a filament nucleation property for CBgs-ParR (Fig. 1E). This is similar to observations for ParM-R1 (Garner *et al.*, 2007; Gayathri *et al.*, 2012). However, after the initial polymerization, CBgs-ParR destabilized the CBgs-ParM filaments and depolymerization occurred, indicating that CBgs-ParR also acted as a depolymerization factor (Fig. 1E). A sedimentation assay also indicated the destabilization of the ParM filament by ParR after 30 min incubation (Fig. S2).

### CryoEM imaging

Cryo-electron microscopy (CryoEM) images of the CBgs-ParM filaments were obtained under three conditions: (i) CBgs-ParM polymerized with ATP for 30 min, (ii) CBgs-ParM polymerized with GTP for 30 min, and (iii) CBgs-ParM polymerized with GTP for 1 min. Filaments with similar diameters were observed under all three conditions (Figs. 2A-C). However, 2D classification of the images showed substantially different averaged images (Figs. 2D-F). The average images of CBgs-ParM, polymerized with ATP, revealed obvious wide and narrow regions on the filament (Fig. 2D), indicating a possible strand cross over. By contrast, the averaged images of CBgs-ParM polymerized with GTP and a short incubation time, showed a more uniform diameter along the filament (Fig. 2F). Finally, the averaged images for the longer polymerization of CBgs-ParM with GTP appeared to show a mixture of the two states (Fig. 2E).

**Figure 2:**
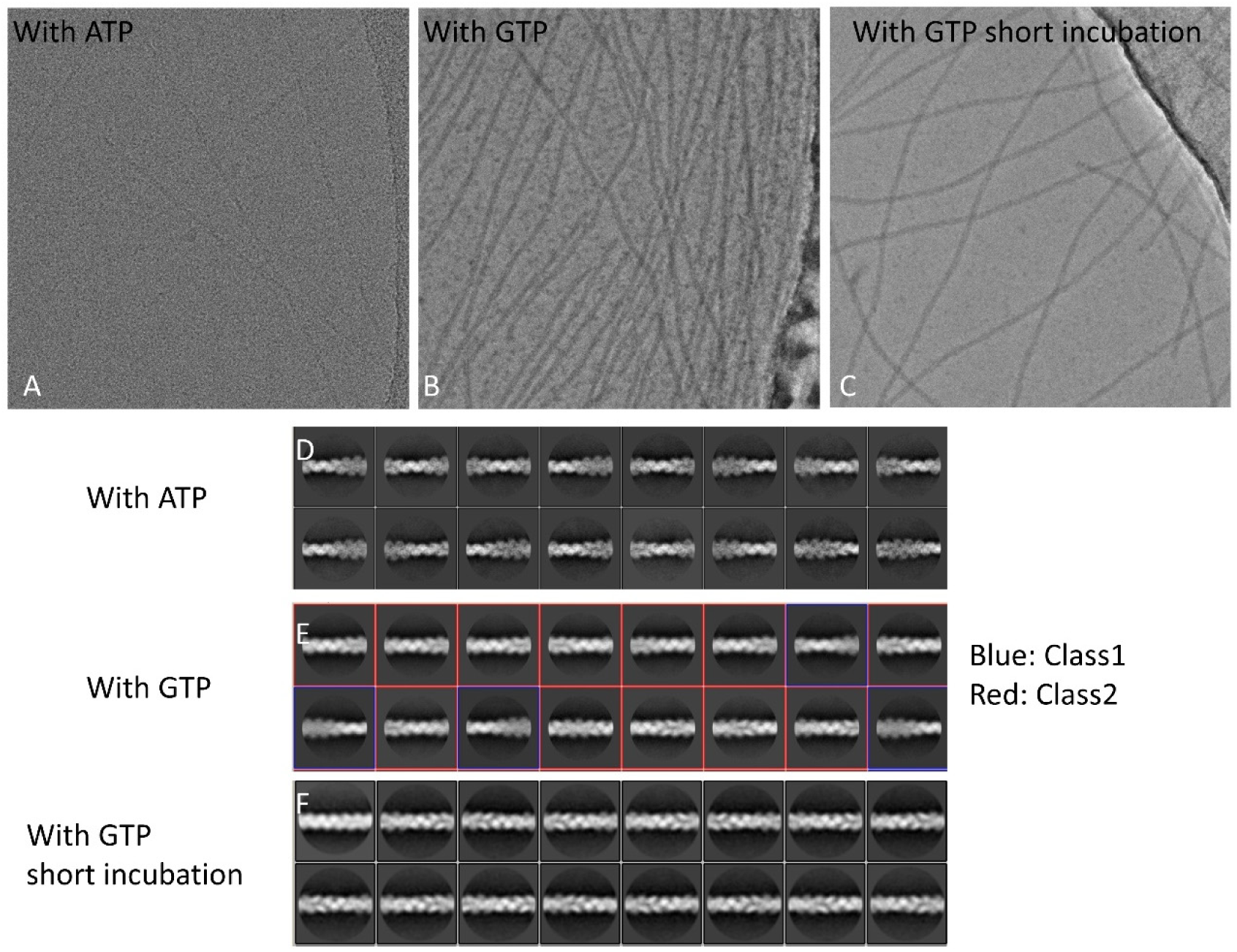
Cryo EM images of CBgs-ParM filaments. A, B, C Cryo EM images of CBgs-ParM filament formed with ATP, with GTP, or with GTP and short incubation time, respectively. D, E, F Averaged images after a 2D classification. The images with GTP were classified into two classes: class1, similar to those with ATP; and class2, similar to those with GTP and short incubation time.

Four density maps were reconstructed for the three polymerization conditions (Fig. 3 and Table S1). One map corresponds to the first condition (ATP, 30 min), at 3.9 Å resolution. Two maps were reconstructed from the second condition (GTP, 30 min), corresponding to the two groups in the 2D classification, class1 and class2 (Fig. 2E). The map from class1 at 3.5 Å resolution was very similar to that from the first condition (ATP, 30 min). The two models from the two maps (ATP 30 min, and class1 with GTP 30 min), were identical up to the reliable resolution of the data, despite the difference in the bound nucleotides. In both maps, density due to gamma phosphate or inorganic phosphate around the nucleotide was not observed, indicating that the bound nucleotides were ADP and GDP, respectively. The maps from class2 with GTP (30 min) and the third condition (GTP, 1 min) were similar to each other, although the resolution was limited, suggesting that they were in the state before the phosphate release.

**Figure 3:**
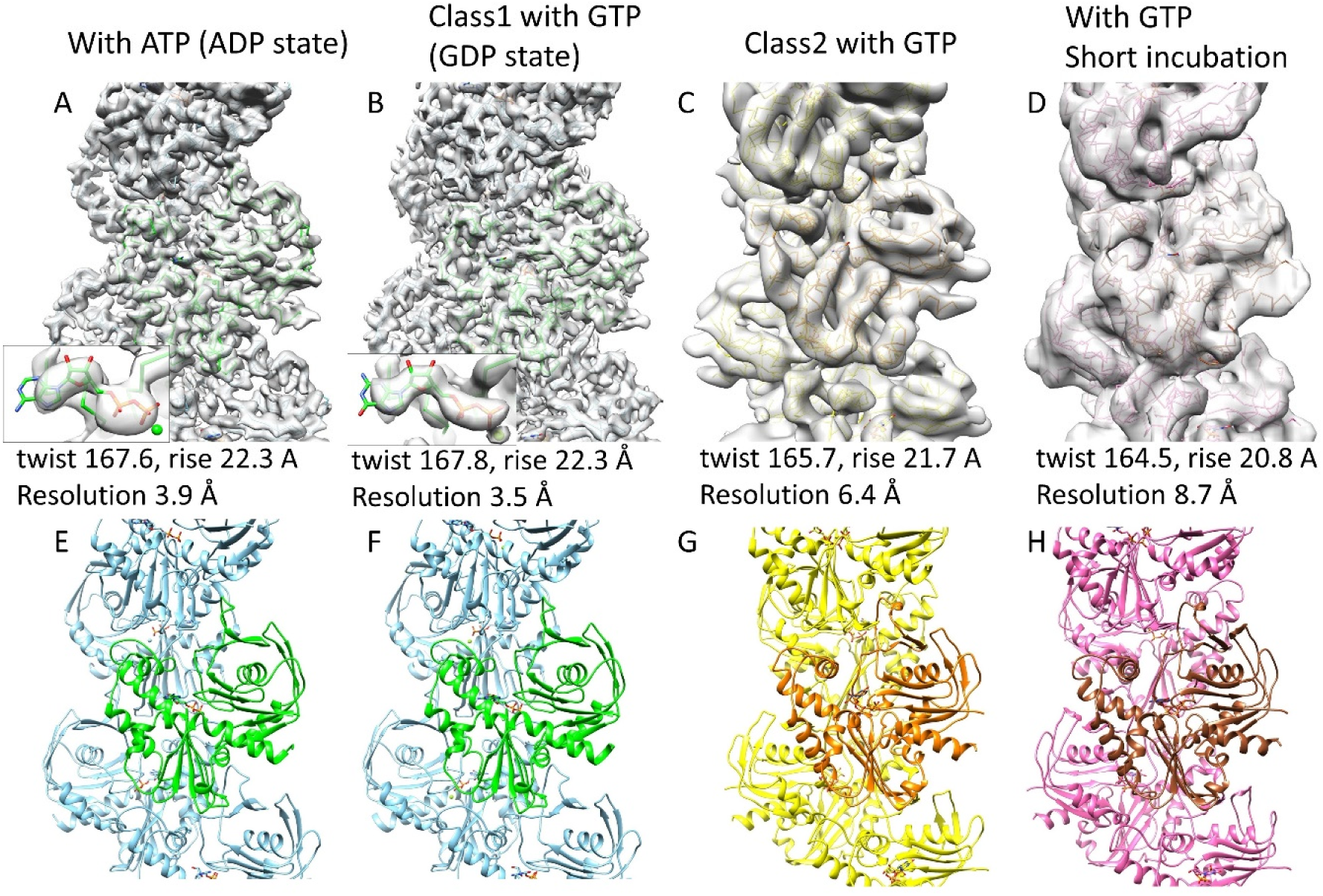
Maps and models of CBgs-ParM structure states with bound nucleotides. A, E with ATP B, F Class1 with GTP C, G Class2 with GTP D, H with GTP and short incubation time The structures with ATP (A and E) were almost identical to that of class1 with GTP (B and F). The gamma phosphate could not be observed in A and B (insets), showing that the binding nucleotides were ADP and GDP, respectively. The model for class2 with GTP (C and G) and the model with GTP and a short incubation time (D and H) were similar to each other.

### Crystal structure, rigid bodies and domain movement

We successfully obtained a crystal structure of the apo form of a mutant of CBgs-ParM, without bound nucleotide (Fig. 4B and Table S2), in which the nucleotide-binding cleft was wide open. This mutant CBgs-ParM was designed to prevent filament assembly via substitution of three residues in the protomer subunit:subunit interface, R204D, K230D and N234D. We compared the high-resolution model built into the cryoEM map of the GDP state filament (Fig. 4A) with the crystal structure of the monomer without nucleotide. We identified two regions that remained identical in the large structural change between the models, similar to those in actin (Oda *et al*, 2019; Tanaka *et al*, 2018). We named these regions, the inner domain (ID) rigid body and the outer domain (OD) rigid body (Figs 4A and B), after actin rigid bodies. No hydrogen bonds were found between the region 144-266, almost the same as the ID rigid body, and the rest of the protein in either state, indicating that the ID rigid body can move independently from the rest of the protein. In the GDP state, instead of direct interactions between residues, GDP and a Mg^2+^ connect the two rigid bodies via hydrogen bonds and salt bridges (Fig 4C). This explains why the crystal structure without nucleotides was wide open. In addition, the guanine moiety of the GDP did not form any contacts or hydrogen bonds with the protein, explaining why CBgs-ParM can utilize both GTP and ATP. The density of guanine and adenine moieties was relatively weak, indicating the flexibility of these parts due to their lack of interaction with the protein (Figs. 3A and B). The OD rigid body interacts with GDP via the beta phosphate and Mg^2+^ (Fig. 4C). Thus, it can be considered that the cleft opening is dependent on the nucleotide state because this binding network would be significantly affected by the presence of another phosphate. Because of the limited resolution, we constructed models for the class2 with GTP and the short incubation time with GTP states with constraints keeping the two rigid bodies (Fig. 3G and H). In comparison with the GDP state model, the nucleotide-binding cleft was slightly open in these two states (Fig. 4D). The models of both states were structurally similar, and they most likely correspond to the GTP or the GDP-Pi state before phosphate release. However, the resolution was not sufficient to confirm the nucleotide state.

**Figure 4:**
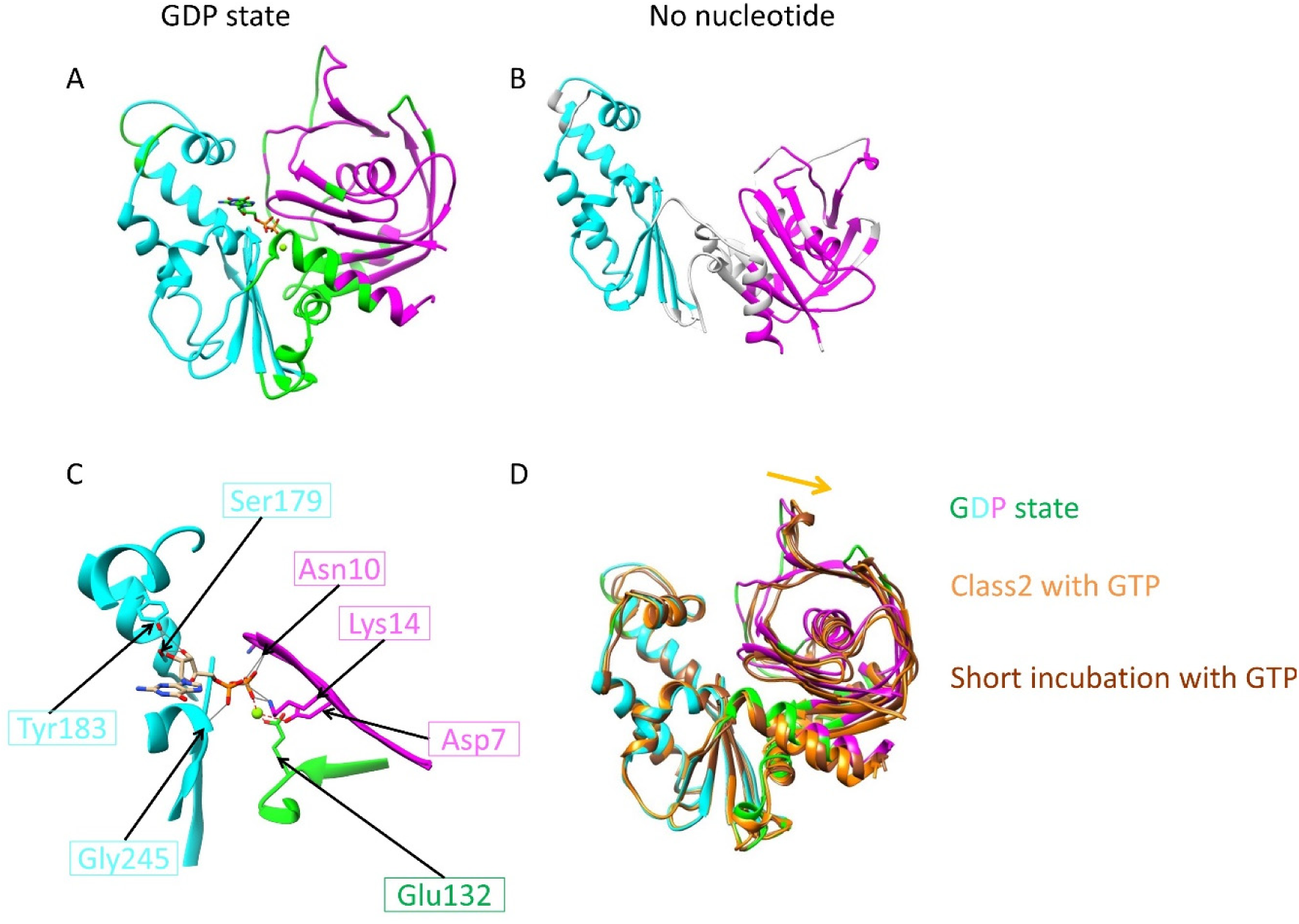
Identification of rigid bodies in CBgs-ParM. A cryo EM model with GDP (Fig. 3B) B crystal structure without nucleotide Two rigid bodies were identified by comparing these structures (ID rigid body, cyan and OD rigid body, magenta). C The bound GDP and Mg^2+^ connected the two rigid bodies (ID rigid body in cyan, OD rigid body in magenta and the rest of the protein in green). Possible hydrogen bonds corresponding to GDP were determined using UCSF chimera (Pettersen *et al*, 2004) and presented as gray lines and possible salt bridges with the Mg^2+^ presented by red dotted lines, although additional hydrogen bonds via water molecules may exist. D Models for the class2 with GTP (orange) and the short incubation time with GTP (brown) were aligned by the ID rigid body and superposed on the model with GDP (green, cyan, and magenta). The nucleotide binding cleft was open in the models for the class2 with GTP and short incubation time with GTP.

### Filament structure

The model with ATP was identical to that of class1 with GTP. Therefore, we compared the three remaining filament models: (i) class1 with GTP (GDP state), (ii) class2 with GTP, and (iii) short incubation time with GTP. The intra-strand interactions between subunits were very similar in the three models (Fig. 5A and B), except for a slight shift in the position of the adjacent subunit in the GDP state. This difference can be explained by the closure of the cleft in the GDP state, which can push the upper subunit leftwards. Despite the similarity within single stands, a large change was observed between strands in forming the filament structure. When the ID rigid body of one subunit of each state was aligned with each other, the position of opposite strand was significantly different (Fig. 5C-D, Movie S1), with a shift as large as 2.5 nm (Figs 5D and E). More numerous inter-strand contacts were found in the GDP state (Fig. 5F) compared to the class2 state with GTP (Fig. 5G), suggesting stabilization of the filament bound to GDP.

**Figure 5:**
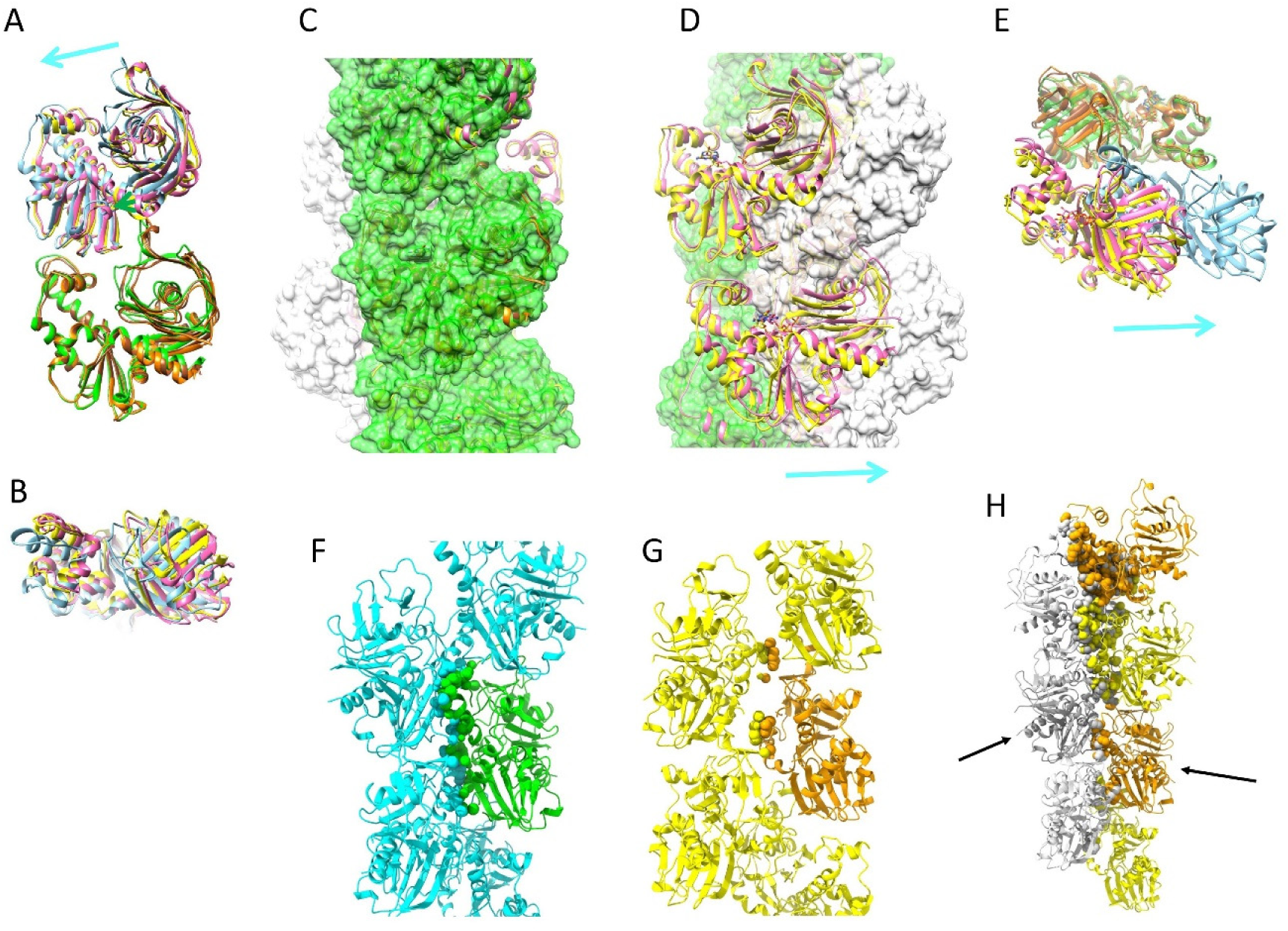
Structural shift in ParM filaments. A, B Two subunits in one strand (A: front view and B: top view). The ID rigid bodies of the lower subunit were aligned with each other. GDP state: green and cyan, Class2 with GTP: orange and yellow, short incubation with GTP: brown and pink. The interactions between the subunits appear similar in the three states except for the upper subunit position, which was slightly shifted in the GDP state compared to the other two states. The closure of the cleft in the GDP state may explain this difference because the closure of the cleft can push the upper subunit leftward. A cyan arrow indicates the direction of the shift of the upper subunit. C-E Relative positions of the two strands. C The model with GDP is presented as a surface model in green and white. Ribbon models for the class2 with GTP (orange and yellow) and the short incubation time with GTP (brown and pink) were superposed. The ID rigid bodies of the center subunits were aligned with each other. D A 180 ° rotated view of Fig. 5C. The opposite strand position in the GDP state was completely different from that in the other two states (compare grey surface to cartoon). E Top view of two adjacent subunits in the different strands. GDP state: green and cyan, Class2 with GTP: orange and yellow, Short incubation with GTP: brown and pink. The ID rigid bodies of the lower subunit (green, orange and brown) were aligned with each other. A cyan arrow indicates the direction of the shift in D and E. F Inter-strand interactions in the CBgs-ParM filament model with GDP. Atoms from the center molecule (green) in contact with the other strand (cyan) were presented in space filling models. 176 contacts were found. G Inter-strand interactions in the model for the class2 with GTP. 36 contacts were found. H The strands of the model with GDP were replaced by the strands of the model for the class2 with GTP. The strands were aligned by the molecules indicated by black arrows. The clashing atoms were presented in space filling models. The contacts and clashes were identified by ChimeraX (Goddard *et al.*, 2018).

### Comparison with CBH-ParM

We compared the CBgs-ParM filament structures with CBH-ParM filament structure, another ParM from *Clostridium botulinum* (Koh *et al.*, 2019). The core region was almost identical. However, the regions contributing to the subunit-subunit interactions largely differed (Figure 6), which leads to differences in the rotational angles and the distances between the subunits in the two strands.

**Figure 6.**
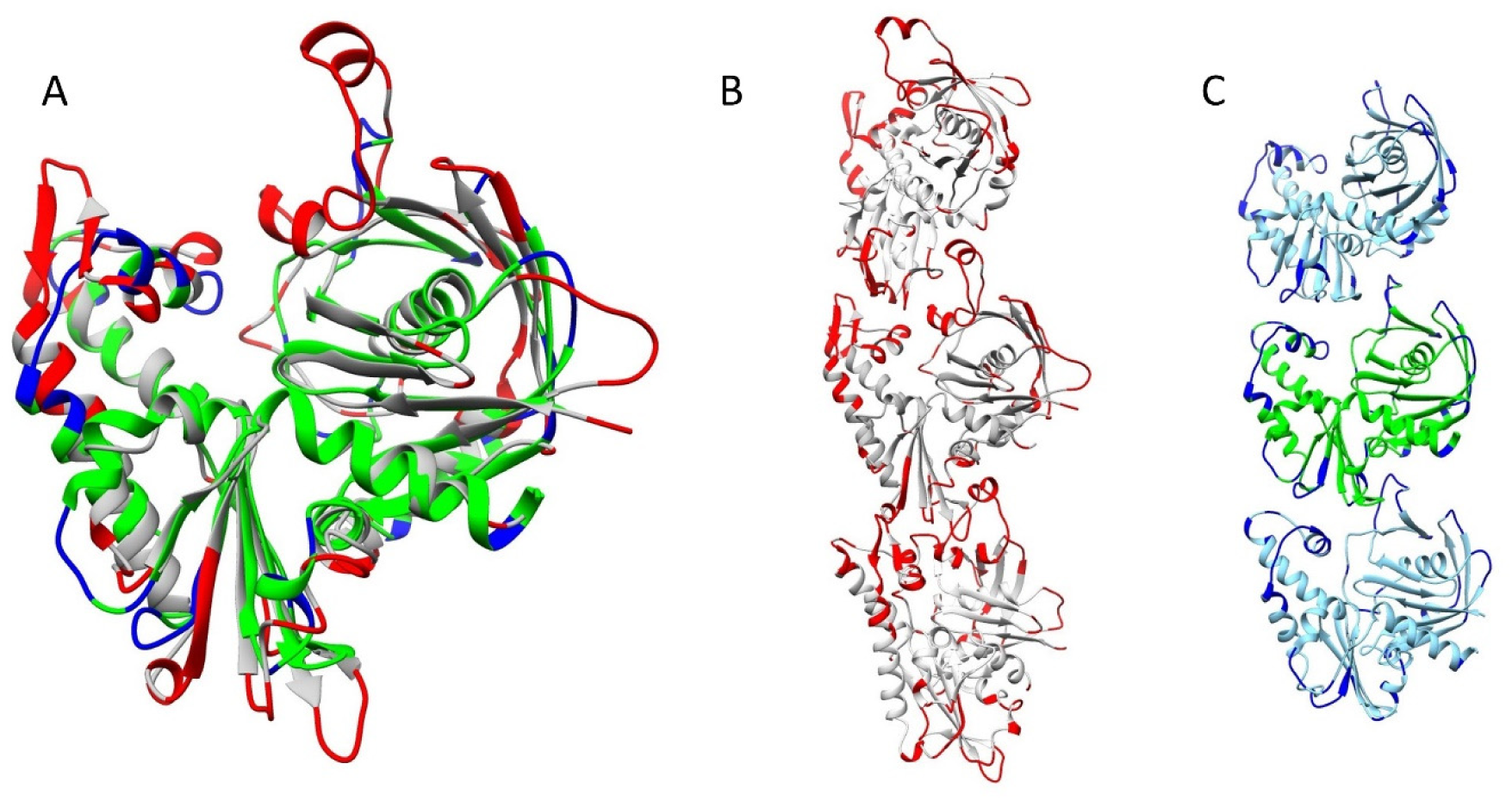
Comparison with CBH-ParM. A Comparison of subunit structures. CBgs-ParM with GDP (green, cyan and blue) and CBH-ParM (gray and red) subunits were structurally aligned by UCSF Chimera (Pettersen *et al.*, 2004). The residues that are more than 3 Å apart from the corresponding residue or that does not have corresponding residue were colored in blue (CBgs-ParM) and red (CBH-ParM). B CBH-ParM strand. Molecular distance and twist in the strand are 5.03 nm and 50.4 degrees, respectively. C CBgs-ParM strand with GDP. Molecular distance and twist in the strand are 4.64 nm and −24.4 degrees, respectively.

## Discussion

CBgs-ParMRC was identified from a whole-genome shotgun sequence and we named it as CBgs-ParMRC to reflect the source. However, there is a possibility that this cassette originates from either a plasmid or from the genome, due to contamination from plasmids during genome isolation. We found gene clusters with high similarity to the sequence around CBgs-ParMRC in the genomes of other *Clostridium* strains, but not in plasmids (Figs. 7 and S3). Therefore, at the current level of genome sequencing we conclude that CBgs-ParMRC is probably encoded in the genome of a limited number of *Clostridium* strains. We speculate that CBgs-ParMRC might contribute to genome segregation in bacteria, similar to microtubules in eukaryotes, although further investigation is required to elucidate its true function.

**Figure 7.**
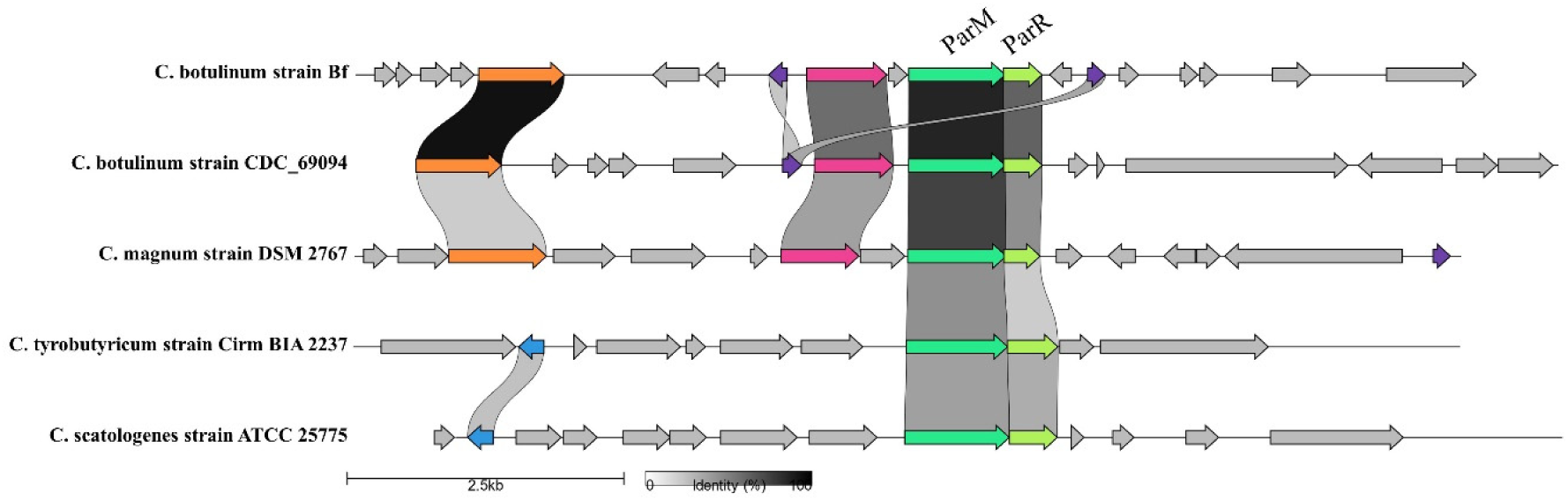
Gene clusters of *Clostridium* species containing ParMRC system. Genes within 5,000 bp of ParM are depicted using clinker&clutermap (Gilchrist & Chooi, 2021). *Clostridium* species containing homologous ParM sequences (identity cutoff = 50 %) are aligned. Genes are depicted as arrows. Conserved genes, ParM (*light green*), ParR (*lime green*), putative replication initiation factor (*magenta*), sporulation-specific N-acetylmuramoyl-L-alanine amidase (*orange*), dihydrodipicolinate reductase (*purple*), and Cro/CI family transcriptional regulator-like protein (*blue*), are colored. Annotations are from GenBank records. *parC* is not displayed because *parC* sequence is not preserved among strains.

The cryoEM images showed obvious differences in the distribution of the nucleotide states between ATP and GTP at 30 mins after initiation of polymerization. It implies either that phosphate release from the ADP-Pi state is faster than that from GDP-Pi state, or faster nucleotide hydrolysis, shorting time in the states of ADP-Pi and ATP in the nucleotide hydrolysis cycle. This is consistent with the faster phosphate release observed with ATP under steady state conditions following the initial polymerization (Fig. 1D, gradient from 10 mins).

Unlike other ParMs, CBgs-ParM polymerizes with ADP or GDP (Fig. 1C) (Garner *et al.*, 2004; Jiang *et al.*, 2016; Koh *et al.*, 2019). The CBgs-ParM filaments, polymerized with either di-phosphate nucleotide, were stable (Fig. S1). Phosphate release from CBgs-ParM polymerized from ATP or GTP did not cause filament instability, as observed for other ParM systems, rather the filament underwent a significant structural change. More inter-strand contacts were observed in the GDP-bound state, providing stability after phosphate release (Fig. 5F and G). When the strands in the GDP model were replaced by the class2 model with GTP, in silico, severe clashes were observed (Fig. 5H). Therefore, the small change in the strand structure (Fig. 5A) caused by the opening of the nucleotide binding cleft (Fig. 4D) in the class2 model with bound GTP prevents the formation of the close contacts observed in the model with GDP. After phosphate release, the strands adopt the GDP conformation allowing for new inter-strand contacts. Thus, phosphate release results in a large structural change in the filament.

We discovered that CBgs-ParM is depolymerize the filaments by a different mechanism to filament instability following phosphate release. CBgs-ParR acts as a nucleating factor during the initial polymerization stage, but later acts as a depolymerization factor for the aged filament. We speculate that ParR recognizes the large structural change in ParM after phosphate release allowing it to change its role to depolymerization. This mechanism has parallels in the eukaryotic actin system, where cofilin senses the nucleotide status of the actin filament, resulting in depolymerization (Carlier *et al*, 1997).

The biochemical and dynamic features of CBgs-ParMR described above are unique among ParMRC systems. However, many ParMRC systems show different features, indicating a large diversity in this family of DNA segregation systems. For instance, another ParM from *C. botulinum* forms a 15-stranded filament with completely different inter-strand interactions (Koh *et al.*, 2019). A ParM from *Bacillus thuringiensis* forms a double-stranded non-polar filament without ParR and a four-stranded filament with ParR, indicating that ParR acts as a template of the filament like gammatubulin for microtubules (Jiang *et al.*, 2016). The *Clostridium tetani* ParM forms a four-stranded filament (Popp *et al.*, 2012), in contrast to the ParM from the *E. coli* R1 plasmid, which forms a double-stranded polar filament similar to the actin filament (Gayathri *et al.*, 2012; Popp *et al*, 2008). In summary, the ParMRC systems form ParM filaments of wide diversity of architectures and dynamics. Usually, a specific ParMRC system is encoded on the plasmid to be segregated. However, the results from the current study indicate that the ParMRC system may also exist on genomes and perhaps contribute to genome segregation. We have previously speculated that when two or more ParMRC systems exist in the same cell, faithful DNA segregation will rely on the two ParMRC systems being orthogonal to each other (Gunning *et al*, 2015). Therefore, a selection pressure to diversify the ParMRC systems may exist to prevent interference between different ParMRC systems, resulting in different architectures of their filament systems. This idea is consistent with our comparison of the ParMs from *Clostridium botulinum* (Fig. 6). The interacting regions between the subunits show larger diversity than the other regions, preventing them from co-polymerization.

## Materials and Methods

### Protein Expression and Purification

The CBgs-ParMRC operon was originally identified through whole-genome shotgun sequencing (ABDP01000001.1) of a clinically isolated strain, *C. botulinum* Bf, which causes infant botulism. Constructs of *parM* and *parR* were synthesized and cloned into pSY5 and pSNAP vectors encoding an 8-histidine tag followed by a human rhinovirus HRV 3C protease cleavage site, respectively. Plasmids were transformed into BL21 (DE3) cells grown to OD600 ~ 0.8, and protein expression was induced with 0.2–1.0 mM IPTG overnight at 15 °C. The cultures were then centrifuged at 4000 × *g* and the cell pellets were resuspended in 50 mM Tris-HCl pH 8.0, 500 mM NaCl, 20 mM imidazole, 5% glycerol, 0.5 mg/mL lysozyme, 0.1 % Triton-X, and protease inhibitor tablets (1 per 2 L culture, Roche, Basel, Switzerland) and lysed via sonication. The cell lysate was then clarified by centrifugation at 30,000 × *g* for 30 min and filtered through a 0.45 μm membrane. The filtered supernatant was loaded onto a HisTrap FF 5 mL (GE Healthcare, Marlborough, MA, USA) preequilibrated with 50 mM Tris-HCl (pH 8.0) containing 500 mM NaCl and 20 mM imidazole. Following a washing step, human rhinovirus HRV 3C protease (5 mg/ml) was loaded in the same buffer for cleavage of tagged proteins (12 h at 4 °C). The cleaved protein was then eluted with washing buffer, pooled, concentrated and subjected to size-exclusion chromatography (Superdex 75 pg, GE Healthcare) in 40 mM HEPES pH 7.5, 150 mM KCl, 2 mM MgCl2, and 1 mM DTT. Fractions were checked for purity via SDS–PAGE, and the pure fractions were pooled and concentrated to between 5 and 10 mg/mL, as determined by UV absorbance at 280 nm using an estimated A280 value calculated using PROTEINCALCULATOR v3.4 (http://protcalc.sourceforge.net).

### Electrophoretic mobility shift assay

The reaction mixture (10 μL) containing 20 nM to 20 μM of ParR in 25 mM HEPES-HCl (pH 7.5), 300 mM KCl, 1 mM MgCl2, 0.5 mM DTT, 1 mg/ml bovine serum albumin, 0.1 μg/μl sonicated salmon sperm DNA, and 5 % glycerol was mixed at 25 °C for 10 min, followed by the addition of 20 nM 5’-FAM-labelled parC DNA fragments and further incubation for 20 min. The polyacrylamide gels were prerun at 150 V for 1 h. After incubation, reactions were analyzed by electrophoresis on a 1 x TBE (pH 7.5), 4% polyacrylamide gel in 1 x TBE running buffer (0.89 M Tris-base, 0.89 M boric acid, 0.02 M EDTA, pH 8.3) at 150 V for 1 h. Gels were scanned using a Pharos FX Plus Molecular Imager (Bio-Rad, Hercules, CA, USA) with an attached external laser.

### Polymerization assays

Assembly and disassembly of CBgs-ParM at 24 °C was followed by light scattering at 90 ° using a Perkin Elmer LS 55 spectrometer for extended time measurements (initial delay time due to mixing by hand ~ 10 s) or a BioLogic stopped-flow machine to observe the early polymerization phase (initial delay time ~ 3 ms), monitored at 600 nm. Assembly was initiated by addition of 2 mM nucleotide at 24 °C in 20 mM Hepes pH 7.5, 350 mM KCl, 2 mM MgCl2.

### Pi release assay

21 μM CBgsParM protein was mixed with the appropriate nucleotide in the same buffer as cryoEM conditions and incubated according to the time course. Reaction mixture was stopped by adding equal volume of ice-chilled 0.4 M perchloric acid (PCA). The reaction mixture was then centrifuged at 1,700 x g for 1 minute. Equal volumes of supernatant and Malachite Green reagent were mixed and incubated at 25 °C for 25 minutes (Kodama *et al*, 1986). Absorbance was read at 650 nm with a Hitachi U-3010 spectrophotometer.

### Sedimentation Assay

To investigate the critical concentrations for polymerization, polymerization of different concentrations of CBgs-ParM (4–15 μM) was initiated by the addition of 5 mM nucleotide (ATP, ADP, GTP, and GDP) in 40 mM HEPES (pH 7.5), 300 mM KCl, 2 mM MgCl2, and 0.5 mM DTT at 24 °C for 30 min. Samples were centrifuged at 279,000 × *g* for 20 min and pellets were resuspended in the same volume as the reaction. Concentrations of CBgs-ParM in the supernatant were estimated via SDS–PAGE, and gel images were analyzed using ImageJ software. The concentrations were not dependent on the total concentration of CBgs-ParM, indicating that they reflected the critical concentration for each nucleotide state. Therefore, the concentrations of the supernatants from different total ParM concentrations were averaged.

To investigate the effects of CBgs-ParR on CBgs-ParM, polymerization of CBgs-ParM (20 μM) with and without CBgs-ParR (20 μM) was initiated by the addition of 5 mM nucleotide (ATP, ADP, GTP, and GDP) in 40 mM HEPES (pH 7.5), 300 mM KCl, 2 mM MgCl2, and 0.5 mM DTT at 24 °C for 30 min. Samples were centrifuged at 279,000 × *g* for 20 min and pellets were resuspended in the same volume as the reaction and the concentrations of CBgs-ParM in the supernatant were estimated via SDS–PAGE.

### Crystallography

CBgs-ParM mutant was constructed with three mutations (R204D, K230D and N234D) to prevent polymerization and allow for crystallization. Purified protein was subjected to crystallization trials by mixing and incubating 5 mg/ml CBgs-ParM mutant and 1 mM AMPPNP on ice for 1 hour. Via the hanging drop vapour diffusion method crystals were grown in 0.5 μl of protein/AMPPNP and 1 μl of mother liquor (0.2 M ammonium chloride, 22% (w/v) PEG 3350) at 288 K. X-ray diffraction data were collected on a RAYONIX MX-300 HS CCD detector on beamline TPS 05A (NSRRC, Taiwan, ROC) controlled by BLU-ICE (version 5.1) at λ= 1.0 Å. Indexing, scaling, and merging of data was performed using HKL2000 (version 715)(Otwinowski & Minor, 1997). Molecular replacement using the protomer built into the 3.5 Å cryoEM density map was carried out in the Phaser (Adams *et al*, 2011) after splitting the structure into the two domains. Model building was carried out in Coot (Emsley & Cowtan, 2004) and refinement in PHENIX (Adams *et al.*, 2011). Data collection and final refinement statistics are summarized in Table S1. Although CBgs-ParM was crystallized in the presence of AMPPNP, the resultant structure did not contain nucleotide.

### Cryo-electron microscopy

CBgs-ParM (0.7 mg/mL) was polymerized in 20 mM HEPES-HCl pH 7.5 containing 250 mM KCl, 1.7 mM MgCl2, and 3 mM GTP or ATP. The mixed solution was incubated for 30 min or 1 min at 25 °C. R1.2/1.3 Mo400 grids (Quantifoil, Jena, Germany) were glow discharged and used within an hour. The reaction mixture (2.5 μL) was applied on the glow discharged grids, blotted on the EM GP (Leica, Wetzlar, Germany) and vitrified by plunging in liquid ethane cooled by liquid nitrogen. Frozen grids were kept under liquid nitrogen for no more than 1 week before imaging. To screen for optimum conditions for cryoEM imaging, the grids were manually observed in a Tecnai G2 Polara (FEI, Hillsboro, OR, USA) cryo transmission electron microscope (at Nagoya University) equipped with a field emission gun operated at 300 kV and a minimal dose system. Images were captured at a nominal magnification of × 115,000 with an underfocus ranging from 1.5 to 3.5 μm and by subjecting the sample to a 2 s exposure time corresponding to an electron dose of ~30 electrons per Å^2^. Images were recorded on a GATAN US4000 CCD camera using an energy filter operated between 10 and 15 eV, with each pixel representing 1.8 Å at the specimen level at exposure settings. Samples were imaged using a Titan Krios microscope operated at 300 kV installed with EPU software (Thermo Fisher, Waltham, MA, USA) at Osaka University. The imaging parameters were actual defocus 1.0–3.0 μm, dose rate 45 e^-^/Å^2^/s, exposure time 1 s, and three image acquisitions per hole. The images with ATP were recorded with a Falcon II detector (Thermo Fisher) at a pixel size of 0.87 Å/pixel with an objective aperture 100 μm, while the images with GTP and 30 min incubation time were recorded with a Falcon III (Thermo Fisher) at a pixel size of 0.87 Å/pixel with an objective aperture 100 μm. The images with GTP and 1 min incubation were recorded with a Falcon III (Thermo Fisher) at a pixel size of 0.87 Å/pixel with a phase plate.

### Image processing

From the sample with ATP, 2,778 images were collected. Image processing was performed using RELION 3.1 (He & Scheres, 2017; Scheres, 2012) software. After motion correction and contrast transfer function (CTF) estimation with CTFFIND-4.1 (Rohou & Grigorieff, 2015), 1,868 images were selected for further image processing. Filaments were manually picked with e2helixboxer, after which particles were extracted at a box size of 384 × 384 pixels. After 2D classification, 36,762 particles were selected. The initial 3D reference was prepared using conventional helical reconstruction using EOS (Yasunaga & Wakabayashi, 1996). Helical symmetry converged to 167.6° twist/22.3 Å rise along the left-handed helix, and the resolution reached 3.9 Å. With GTP and 30 min incubation time, 1,398 images were collected. After motion correction and CTF estimation, 152,490 particles were extracted. We categorized Class2D averaged images into two by visual inspection. The first category contained 40,599 particles and the helical symmetry converged to 167.8 ° twist/22.3 Å at 3.5 Å resolution, while the second contained 70,754 particles and helical symmetry converged to 165.7° twist/21.7 Å rise at 6.5 Å resolution. For the map with GTP and 1 min incubation time, 2,772 images were collected and 153,326 particles were used for the final reconstruction at 8.6 Å resolution.

### Model building

The initial atomic model with GDP was constructed by homology modeling using Rosetta3 with the pCBH ParM model (Koh *et al.*, 2019) (6IZV) as a template. The resulting model was iteratively refined using COOT (Emsley *et al*, 2010), molecular dynamics flexible fitting (MDFF, using ISOLDE (Croll, 2018), an extension of ChimeraX (Goddard *et al*, 2018)), and Phenix (Adams *et al*, 2010). GDP in the final GDP model was replaced by ADP to give the initial model with ADP, which was then refined using the same procedures. The final GDP model was also used as the initial model for the lower resolution structures (PDBIDs 7X55 and 7X59), which were fitted into the map by MDFF with adaptive distance restraints for the two rigid bodies using ISOLDE(Croll, 2018). The resultant model was refined using COOT and Phenix.

### Rigid body search

The model with GDP and the crystal structure without nucleotides were aligned with each other to maximize the number of Cα with less than 0.7 Å deviation between the two models. The resultant residues with less than 0.7 Å deviation were considered as the rigid body(Tanaka *et al.*, 2018). Two rigid bodies were identified (Figure 4).

**Figure 8:**
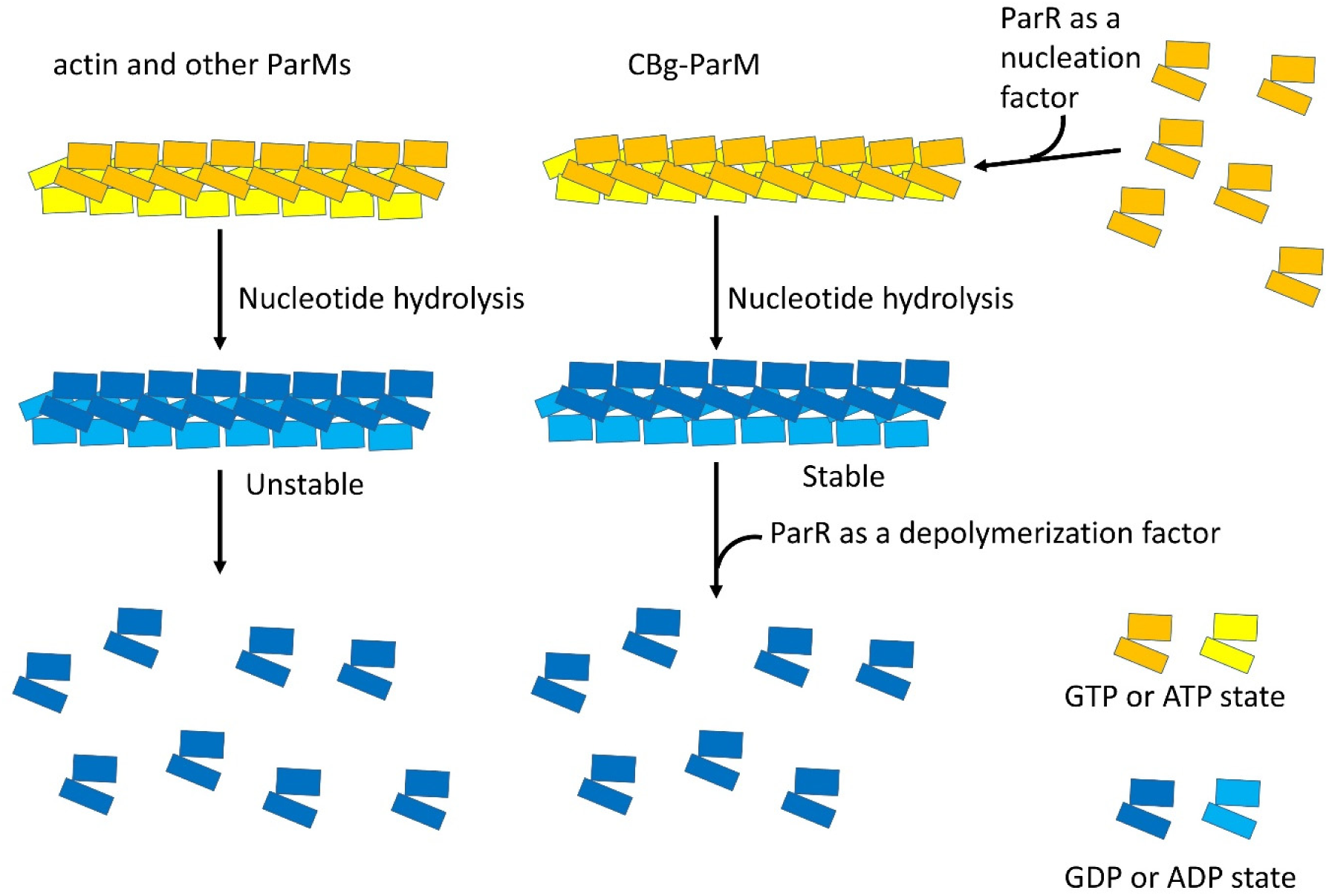
Polymerization and depolymerization of bacterial actin homologs. Usual ParMs, form a stable filament with ATP or GTP. ATP or GTP is hydrolyzed in the filament, which makes the filament unstable, resulting in depolymerization. CBgs-ParM has different features, with ATP/GTP hydrolysis accompanied by a significant change in the filament structure without large destabilization. ParR acts as a depolymerizing factor for CBgs-ParM with ADP or GDP, in addition to a nucleation activity for CBgs-ParM with ATP and GTP.

## Data availability

The CBgs-ParM filament coordinates from this publication have been deposited in the Protein Data Bank (PDB, https://www.rcsb.org/) under accession codes 7X54, 7X56, 7X59 and 7X55 for ADP state, GDP state, the second class with GTP, and the short incubation time with GTP, respectively. The corresponding EM maps have been deposited in the Electron Microscopy Data Bank (EMDB, https://www.ebi.ac.uk/emdb/) (EMD-33007, 33009, 33012 and 33008). The apo CBgs-ParM X-ray structure is deposited in the PDB (7X3H). All other data are available from the corresponding author upon reasonable request.

## Acknowledgments

This research was supported by JSPS KAKENHI (grant numbers 18H02410 and 21H02440 to AN), JST CREST (JPMJCR19S5 to RCR and AN), and by Vidyasirimedhi Institute of Science and Technology (VISTEC, RCR). This research was also supported by Nanotechnology Platform Program of the Ministry of Education, Culture, Sports, Science and Technology (MEXT), Japan, Grant Number JPMXP09A19OS0052 at the Research Center for Ultra-High Voltage Electron Microscopy (Nanotechnology Open Facilities) in Osaka University and by the Collaborative Research Program of Institute for Protein Research, Osaka University, CEMCR-1701.

## Author Contributions

AK, RCR, and AN designed the study. AK, SA, and DP performed protein purification. SA and DP performed biochemical experiments and light microscopy. AK, NM, KI, and AN performed electron microscopy. RCR solved the crystal structure. AK and AN performed image analysis and model building. RCR performed crystallography. YK performed sequence analysis. AK, SA, RCR, and AN wrote the manuscript.

## References

Adams PD, Afonine PV, Bunkoczi G, Chen VB, Davis IW, Echols N, Headd JJ, Hung LW, Kapral GJ, Grosse-Kunstleve RW et al (2010) PHENIX: a comprehensive Python-based system for macromolecular structure solution. Acta crystallographica Section D, Biological crystallography 66: 213–221

Adams PD, Afonine PV, Bunkoczi G, Chen VB, Echols N, Headd JJ, Hung LW, Jain S, Kapral GJ, Grosse Kunstleve RW et al (2011) The Phenix software for automated determination of macromolecular structures. Methods 55: 94–106

Bharat TA, Murshudov GN, Sachse C, Lowe J (2015) Structures of actin-like ParM filaments show architecture of plasmid-segregating spindles. Nature 523: 106–110

Carlier MF, Laurent V, Santolini J, Melki R, Didry D, Xia GX, Hong Y, Chua NH, Pantaloni D (1997) Actin depolymerizing factor (ADF/cofilin) enhances the rate of filament turnover: implication in actin-based motility. The Journal of cell biology 136: 1307–1322

Croll TI (2018) ISOLDE: a physically realistic environment for model building into low-resolution electron-density maps. Acta crystallographica Section D, Structural biology 74: 519–530

Emsley P, Cowtan K (2004) Coot: model-building tools for molecular graphics. Acta crystallographica Section D, Biological crystallography 60: 2126–2132

Emsley P, Lohkamp B, Scott WG, Cowtan K (2010) Features and development of Coot. Acta crystallographica Section D, Biological crystallography 66: 486–501

Fujiwara I, Vavylonis D, Pollard TD (2007) Polymerization kinetics of ADP-and ADP-Pi-actin determined by fluorescence microscopy. Proceedings of the National Academy of Sciences of the United States of America 104: 8827–8832

Garner EC, Campbell CS, Mullins RD (2004) Dynamic instability in a DNA-segregating prokaryotic actin homolog. Science 306: 1021–1025

Garner EC, Campbell CS, Weibel DB, Mullins RD (2007) Reconstitution of DNA segregation driven by assembly of a prokaryotic actin homolog. Science 315: 1270–1274

Gayathri P, Fujii T, Moller-Jensen J, van den Ent F, Namba K, Lowe J (2012) A bipolar spindle of antiparallel ParM filaments drives bacterial plasmid segregation. Science 338: 1334–1337

Gilchrist CLM, Chooi YH (2021) Clinker & clustermap.js: Automatic generation of gene cluster comparison figures. Bioinformatics

Goddard TD, Huang CC, Meng EC, Pettersen EF, Couch GS, Morris JH, Ferrin TE (2018) UCSF ChimeraX: Meeting modern challenges in visualization and analysis. Protein science: a publication of the Protein Society 27: 14–25

Gunning PW, Ghoshdastider U, Whitaker S, Popp D, Robinson RC (2015) The evolution of compositionally and functionally distinct actin filaments. J Cell Sci 128: 2009–2019

He S, Scheres SHW (2017) Helical reconstruction in RELION. Journal of structural biology 198: 163–176

Jiang S, Narita A, Popp D, Ghoshdastider U, Lee LJ, Srinivasan R, Balasubramanian MK, Oda T, Koh F, Larsson M et al (2016) Novel actin filaments from Bacillus thuringiensis form nanotubules for plasmid DNA segregation. Proceedings of the National Academy of Sciences of the United States of America 113: E1200–1205

Kodama T, Fukui K, Kometani K (1986) The initial phosphate burst in ATP hydrolysis by myosin and subfragment-1 as studied by a modified malachite green method for determination of inorganic phosphate. Journal of biochemistry 99: 1465–1472

Koh F, Narita A, Lee LJ, Tanaka K, Tan YZ, Dandey VP, Popp D, Robinson RC (2019) The structure of a 15-stranded actin-like filament from Clostridium botulinum. Nature communications 10: 2856

Narita A (2011) Minimum requirements for the actin-like treadmilling motor system. Bioarchitecture 1: 205–208

Oda T, Takeda S, Narita A, Maeda Y (2019) Structural Polymorphism of Actin. Journal of molecular biology 431: 3217–3228

Otwinowski Z, Minor W (1997) Processing of X-ray diffraction data collected in oscillation mode. Methods in enzymology 276: 307–326

Pettersen EF, Goddard TD, Huang CC, Couch GS, Greenblatt DM, Meng EC, Ferrin TE (2004) UCSF Chimera--a visualization system for exploratory research and analysis. Journal of computational chemistry 25: 1605–1612

Pollard TD, Borisy GG (2003) Cellular motility driven by assembly and disassembly of actin filaments. Cell 112: 453–465

Popp D, Narita A, Lee LJ, Ghoshdastider U, Xue B, Srinivasan R, Balasubramanian MK, Tanaka T, Robinson RC (2012) Novel actin-like filament structure from Clostridium tetani. The Journal of biological chemistry 287: 21121–21129

Popp D, Narita A, Oda T, Fujisawa T, Matsuo H, Nitanai Y, Iwasa M, Maeda K, Onishi H, Maeda Y (2008) Molecular structure of the ParM polymer and the mechanism leading to its nucleotide-driven dynamic instability. The EMBO journal 27: 570–579

Rohou A, Grigorieff N (2015) CTFFIND4: Fast and accurate defocus estimation from electron micrographs. Journal of structural biology 192: 216–221

Salje J, Gayathri P, Lowe J (2010) The ParMRC system: molecular mechanisms of plasmid segregation by actin-like filaments. Nature reviews Microbiology 8: 683–692

Scheres SH (2012) RELION: implementation of a Bayesian approach to cryo-EM structure determination. Journal of structural biology 180: 519–530

Tanaka K, Takeda S, Mitsuoka K, Oda T, Kimura-Sakiyama C, Maeda Y, Narita A (2018) Structural basis for cofilin binding and actin filament disassembly. Nature communications 9: 1860

Wegner A (1976) Head to tail polymerization of actin. Journal of molecular biology 108: 139–150

Yasunaga T, Wakabayashi T (1996) Extensible and object-oriented system Eos supplies a new environment for image analysis of electron micrographs of macromolecules. Journal of structural biology 116: 155–160

