## Supplementary data for "An actin-like filament from *Clostridium botulinum* exhibits a novel mechanism of filament dynamics"

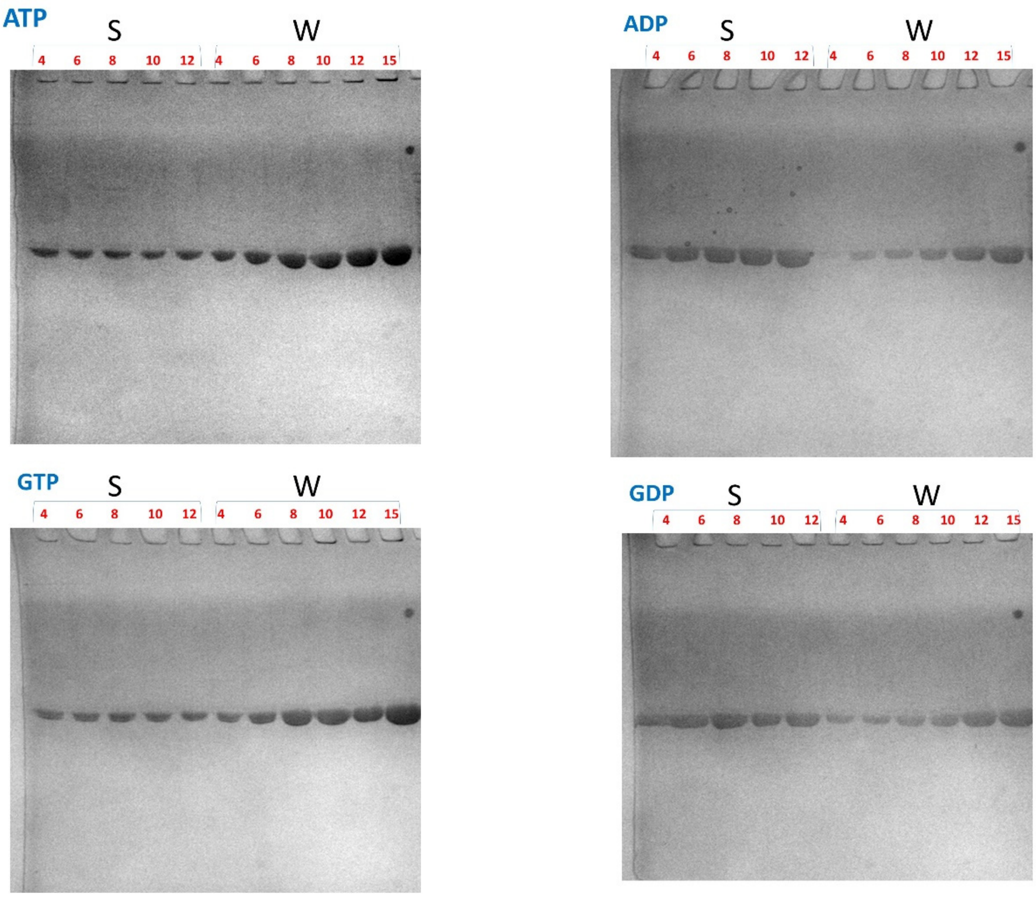

**Figure S1. Sedimentation assay for estimating critical concentrations.** S represents supernatant and W represents whole before sedimentation. The number above each lane represents the concentration of ParM (μM). ParM concentrations from W lanes were used to plot a standard curve to estimate the ParM concentrations of S lanes.

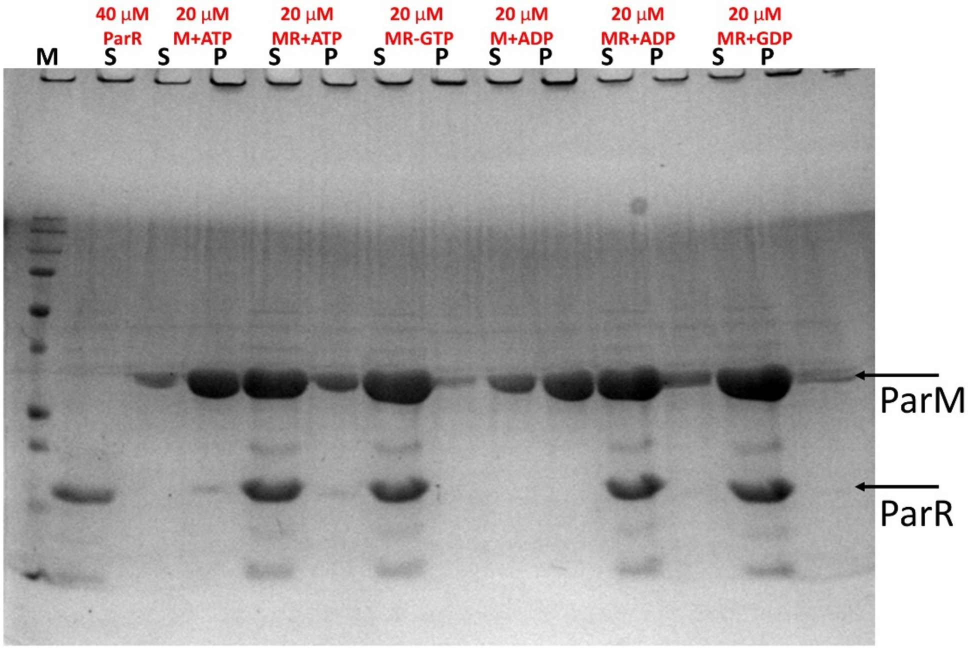

**Figure S2. Sedimentation assay for ParM with and without ParR.** M and MR represent ParM without ParR and with ParR, respectively. S and P represent supernatant and pellet, respectively. The clear decrease in ParM concentration in the pellet fraction with ParR indicates the depolymerizing function of ParR.

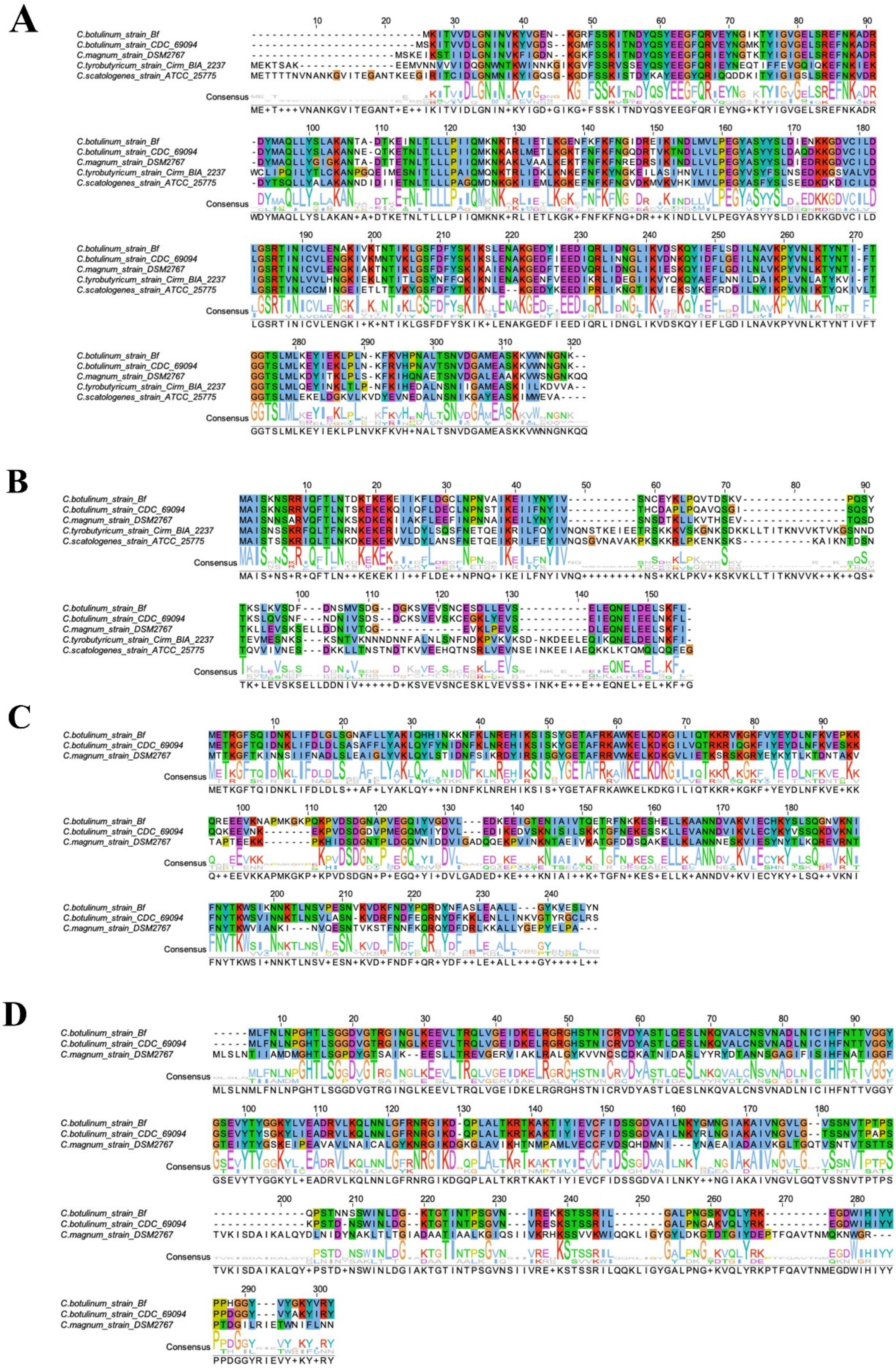

Figure S3: Multiple sequence alignment of homologous sequences of ParMRC gene clusters

**in *Clostridium* sp.** Multiple sequence alignment of ParM (A), ParR (B), putative replication initiator (C), and sporulation-specific N-acetylmuramoyl-L-alanine amidase (D) were performed with MUSCLE (Edgar, 2004). Homologous residues are colored according to the ClustalX scheme.

**Movie S1: Structural shift in ParM filaments.** Models and maps for the class1 (corresponding to GDP state) and the class2 with GTP were aligned by the ID rigid bodies of the center subunit.

**References**

Edgar RC (2004) MUSCLE: multiple sequence alignment with high accuracy and high throughput. *Nucleic Acids Res* 32: 1792-1797

| | #1 name (EMDB-33007) (PDB 7X54) | #2 name (EMDB-33009) (PDB 7X56) | #3 name (EMDB-33012) (PDB 7X59) | #4 name (EMDB-33008) (PDB 7X55) |
|---|---|---|---|---|
| **Data collection and processing** | | | | |
| Voltage (kV) | 300 kV | 300 kV | 300 kV | 300 kV |
| Electron exposure (e–/Å$^2$) | 45 | 45 | 45 | 45 |
| Defocus range (μm) | 1.5~3.5 | 1.0~3.0 | 1.0~3.0 | 0.5~1.5 |
| Pixel size (Å) | 0.87 | 0.87 | 0.87 | 0.87 |
| Phase plate | No | No | No | Yes |
| Symmetry imposed | Helical | Helical | Helical | Helical |
| Final particle images (no.) | 36762 | 40599 | 70754 | 153326 |
| Map resolution (Å)<br>FSC threshold | 3.9<br>0.143 | 3.5<br>0.143 | 6.5<br>0.143 | 8.6<br>0.143 |
| Map resolution range (Å) | ∞ ~ 3.9 | ∞ ~ 3.5 | ∞ ~ 6.5 | ∞ ~ 8.6 |
| **Refinement** | | | | |
| Initial model used (PDB code) | 7X56 | 6IZV | 7X56 | 7X56 |
| Map sharpening *B* factor (Å$^2$) | -105 | -117 | -527 | -1281 |
| Model composition<br>Non-hydrogen atoms<br>Protein residues<br>Ligands | <br>11575<br>1425<br>10 | <br>11580<br>1425<br>10 | <br>11595<br>1425<br>5 | <br>11600<br>1425<br>5 |
| R.m.s. deviations<br>Bond lengths (Å)<br>Bond angles (°) | <br>0.004<br>0.979 | <br>0.005<br>0.965 | <br>0.004<br>1.059 | <br>0.007<br>1.137 |
| Validation<br>MolProbity score<br>Clashscore<br>Poor rotamers (%) | <br>1.98<br>8.79<br>0.39 | <br>1.96<br>7.71<br>0.00 | <br>2.19<br>11.95<br>0 | <br>2.44<br>21.32<br>0 |
| Ramachandran plot<br>Favored (%)<br>Allowed (%)<br>Disallowed (%) | <br>91<br>9<br>0 | <br>90<br>10<br>0 | <br>88<br>12<br>0 | <br>89<br>11<br>0 |

**Table S1. Cryo-EM data collection, refinement and validation statistics**

| | CBgs-ParM (PDB code 7X3H) |
|---|---|
| **Protein** | |
| Accession No. | EDT87363.1 |
| Mutations | R204D, K230D, N234D |
| **Data collection** | |
| Crystal[illegible] | $P2_1$ |
| *a*, *b*, *c* (Å) | 55.6, 51.1, 64.9 |
| $\alpha, \beta, \gamma$ (°) | 90.0, 115.3, 90.0 |
| Wavelength (Å) | 1.0 |
| Resolution (Å)[a] | 50.0-1.7 (1.73-1.70) |
| $R_{merge}$ | 3.0 (47.3) |
| $R_{meas}$ | 3.5 (58.8) |
| $R_{pim}$ | 1.8 (34.4) |
| *I*/[illegible](*I*) | 37.7 (1.8) |
| $CC_{1/2}$ | (0.736) |
| Completeness (%) | 99.3 (94.1) |
| Redundancy | 3.6 (2.5) |
| **Refinement** | |
| Resolution (Å) | 32.0-1.7 (1.76-1.70) |
| No. reflections | 35734 (2896) |
| $R_{work}$ / $R_{free}$ | 19.2/22.4 (27.1/29.9) |
| No. atoms | |
| Protein | 2249 |
| Water | 314 |
| *B* factors | |
| Protein | 25.4 |
| Water | 34.0 |
| r.m.s deviations | |
| Bond lengths (Å) | 0.007 |
| Bond angles (°) | 1.09 |
| Ramachandran Plot | |
| Favoured (%) | 98.5 |
| Outliers (%) | 0 |

**TableS2. X-ray crystallography data collection, refinement and validation statistics**
